# Discovery of NRG1-VII: a novel myeloid-derived class of NRG1 isoforms

**DOI:** 10.1101/2023.02.02.525781

**Authors:** Miguel Ángel Berrocal-Rubio, Yair David Joseph Pawer, Marija Dinevska, Ricardo De Paoli-Iseppi, Samuel S. Widodo, Josie Gleeson, Nadia Rajab, Will De Nardo, Jeannette Hallab, Anran Li, Theo Mantamadiotis, Michael B. Clark, Christine A. Wells

## Abstract

The growth factor Neuregulin-1 (NRG1) has pleiotropic roles in proliferation and differentiation of the stem cell niche in different tissues. It has been implicated in gut, brain and muscle development and repair. Six isoform classes of NRG1 and over 28 protein isoforms have been previously described. Here we report a new class of NRG1, designated NRG1-VII to denote that these NRG1 isoforms arise from a myeloid-specific transcriptional start site (TSS) previously uncharacterized. Long-read sequencing was used to identify eight high-confidence NRG1-VII transcripts. These transcripts presented major structural differences from one another, through the use of cassette exons and alternative stop codons. Expression of NRG1-VII was confirmed in primary human monocytes and tissue resident macrophages and iPSC-derived macrophages. Isoform switching via cassette exon usage and alternate polyadenylation was apparent during monocyte maturation and macrophage differentiation. NRG1-VII is the major class expressed by the myeloid lineage, including tissue-resident macrophages. Analysis of public gene expression data indicates that monocytes and macrophages are a primary source of NRG1, suggesting that NRG1-VII is the most common class of NRG1 in most adult human tissues, except brain. The size and structure of type VII isoforms suggests that they may be more diffusible through tissues than other NRG1 classes. However, the specific roles of type VII variants in tissue homeostasis and repair have not yet been determined.

## Introduction

Neuregulins (NRGs) are a family of highly pleiotropic growth factors derived from four paralogous genes (*NRG1-4*) (Figure 1A). NRGs are typically synthesized as transmembrane pro-peptides that are cleaved by metalloproteases in the extracellular space to form a bioactive peptide with an exposed epidermal growth factor-like (EGF) domain that can bind erythroblastic leukemia viral oncogene homolog (ERBB) receptors. The human *Neuregulin-1* (*NRG1*) locus, on Chromosome 8p12, generates numerous isoforms (Figure 1A, B) which are thought to be tissue-specific and functionally diverse (Falls, 2003). NRG1 has been implicated in the development of multiple tissues by promoting cell division within the stem cell niche and in differentiation trajectories (Yu et al., 2021, Wagner et al., 2007) including progenitor cells in the gut (Jardé et al., 2020), skeletal muscle (Gumà et al., 2010, Cheret et al., 2013) and cardiac cells (Wagner et al., 2007, Kramer et al., 1996), as well as nervous system development (Birchmeier 2009, Newbern et al., 2010). These studies demonstrate NRG1’s key role in organogenesis and the importance of understanding how NRG1 isoforms exert tissue-specific effects to maintain the adult stem cell niche.

**Figure 1.**
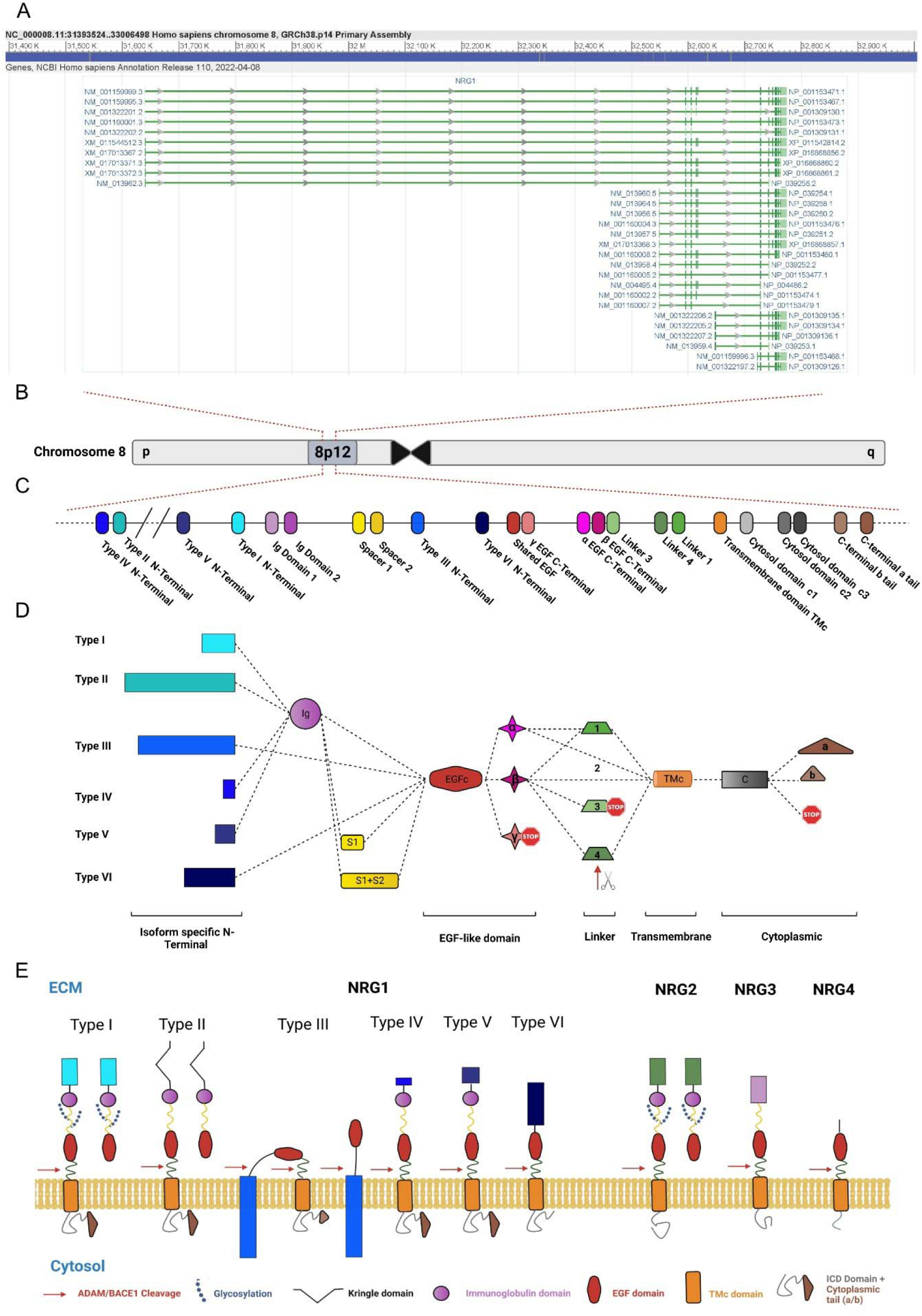
The human *NRG1* locus. **A)** Screenshot of the NCBI RefSeq curated products from the *NRG1* locus showing 28 protein coding transcriptional variants (NCBI Gene website accessed 26/1/2023). Left and right labels correspond to transcript and protein accessions, respectively. **B)** Representation of the NRG1 genomic locus (Chr 8p12). **C)** Schematic annotating exons in the human *NRG1* locus with modular protein coding domains. **D)** Schematic showing combinatorial protein domains for previously annotated NRG1 isoforms. Rows show the six major NRG1 classes, defined by alternate N-terminals. Columns show alternate domains: domains within a column are mutually exclusive. Dotted lines show known connections between domains in different NRG1 isoforms. Red stop symbols represent translation stop. Red arrows represent a protease cleavage point. **E)** Representation of Neuregulin protein isoforms and their domain distribution. Note that in NRG1 types IV, V and VI all isoforms that have been characterized are transmembrane, while I, II and III also present isoforms that don’t. Legend below the cell membrane describes symbols for ADAM/BACE1 cleavage site (red arrow), glycosylation site (blue chain), kringle domain (black V), IgG domain (purple circle), EGF domain (red oval), Transmembrane domain (orange rectangle), intracellular domain (grey squiggle, with or without brown triangle). N-terminal domains of NRG1 reference modularity shown in Fig. 1B and 1C, including the membrane-tethered N-terminal of NRG1 type III.

NRG1 and its receptors are involved in several diseases and targets for clinical research. Germline mutations in NRG1 are associated with developmental brain disorders such as schizophrenia (Craddock et al., 2005) as well as degenerative disorders such as amyotrophic lateral sclerosis (Sun et al., 2020) and Alzheimer’s disease (Go et al., 2005). Constitutively activated isoforms of NRG1 are implicated in cancer, for which blocking the NRG1 isoform Heregulin or its receptors (ERBB) are effective clinical strategies against solid tumours (Sheng et al., 2010, Zhang et al., 2022). Deficiencies in NRG1 are associated with Hirshprung’s disease, leading to poor innervation of the gut (Tang et al., 2012, Garcia-Barcelo et al., 2009), as well as abnormal brain development and mental disorders like bipolar disorder or schizophrenia (Georgieva et al., 2008, Marballi et al., 2012). In contrast, high levels of circulating NRG1 are associated with cardiac disease and morbidity following heart failure (Haller et al., 2022). NRG1 is also involved in modulating the immune response (Alizadeh et al., 2018, Ryzhov et al., 2017), controlling insulin related liver activity (Zhang et al., 2018), cell migration (Jumper et al., 2017) and cell-cell recognition and viability in the central nervous system (Garratt et al., 2000). Therefore, there are clear clinical benefits in understanding which cells express the different NRG1 isoforms in each tissue and how they play their tissue-specific roles in different biological processes.

The diversity of isoform functions in healthy development and disease across different organs is mirrored by the structural diversity and tissue specific transcriptional regulation of NRG1 products. The *NRG1* gene encodes at least 28 isoforms (Figure 1A) according to the ENTREZ Gene reference transcript list (Brown et al., 2015). Others have reported higher isoform heterogeneity from this locus (Mei 2008). This extraordinary transcript diversity is due to the use of alternate transcription initiation sites, cassette exon usage, and alternate polyadenylation sequences. The locus is remarkably modular (Figure 1C), which allows for different structural combinations of NRG1 domains depending on their start site (type I-VI), the use of alternative linker domains, which are the protease targets for ectodomain shedding (1-4), the type of EGF-like domain (α, β or γ), and whether the pro-peptide is membrane tethered or not (Figure 1C-E). One well characterized isoform type, NRG1-III, contains a unique N-terminal domain that includes a sequence that locates to the cell membrane (Nave et al., 2006). This means that upon NRG1 processing, the growth factor domain will not be released from the source cell but will function as a juxtracrine signal that allows neurons to establish key cell-cell interactions (with other neurons, oligodendrocytes, or muscle fibers) and regulates the survival of both interacting cells (Garratt et al., 2000, Wolpowitz et al., 2000). In rare circumstances, another cleavage site between the N-terminal domain and the EGF domain can also be target by a protease, releasing the EGF domain to the ECM (Fleck et al., 2013).

More recently, NRG1 has been described as an important factor regulating the stem cell niche in the gut (Jardé et al., 2020). In this context, NRG1 likely modulates stemness, proliferation and identity of progenitor cells in the niche, and it is required to recapitulate certain secretory and absorptive functions in human gut organoids (Kilik et al., 2021). Beyond development and tissue repair, NRG1 secreted by macrophages has also been implicated in inflammation (Jardé et al., 2020, Garrido-Trigo et al., 2022). However, the specific NRG1 isoforms expressed by macrophages in these contexts have not yet been described.

Determining the sources and types of different NRG1 isoforms is an important part of understanding how NRG1 directs these different developmental, reparative, and inflammatory outcomes. Here, data mining to assess the expression profiles of *NRG1* led to the identification of a previously uncharacterized TSS that appears to be used exclusively in cells of the myeloid lineage. We propose that transcripts generated from this alternative TSS belong to a new NRG1 Class, NRG1-VII. Using Oxford Nanopore sequencing, we identified eight class VII isoforms with distinct transcript structures and predicted the protein characteristics of these isoforms. qRT-PCR targeting the unique first exon of *NRG1-VII* transcripts in human cells confirmed that type VII isoforms are expressed by monocytes, infiltrating macrophages and tissue resident macrophages. Immunohistochemistry using antibodies directed towards the EGF-like or intracellular domains (ICD) demonstrated that tissue-resident macrophages are a major source of NRG1 in these tissues, an observation further supported by transcriptional evidence derived from single cell data collated within the Human Protein Atlas. This study therefore contributes to untangling the complexity of this already intricate locus by characterizing the structure and distribution of myeloid-specific NRG1 isoforms.

## Results

### NRG1-VII is defined by a novel TSS discovered in myeloid cells

Since the different isoforms of NRG1 have varying structures and binding affinities to their receptors, we sought to investigate which of these were expressed by myeloid cells. First, we used the FANTOM5 (Functional Annotation of the Mammalian Genome) database (The FANTOM Consortium; 2014), a catalogue that maps the TSSs of genes expressed in 975 human samples (including cells, tissues, and cell lines). Here, we found a previously uncharacterized TSS of NRG1 that was exclusively active in myeloid cells, including monocytes, macrophages, and basophils (Figure 2A). A schematic of the locus suggested that this represents a potential new class of *NRG1* transcripts which we prospectively named *NRG1-VII* (Figure 2). The FANTOM TSS predicted that the *NRG1-VII* starting exon contained 139 base pairs (bp) of mRNA sequence unique to transcripts originating from this site, and an in-frame methionine was identified 115 bp downstream from the start of transcription (Figure 2B, C). Therefore, we aimed to characterize whether this newly described TSS would lead to the expression of a new class of protein coding transcripts.

**Figure 2.**
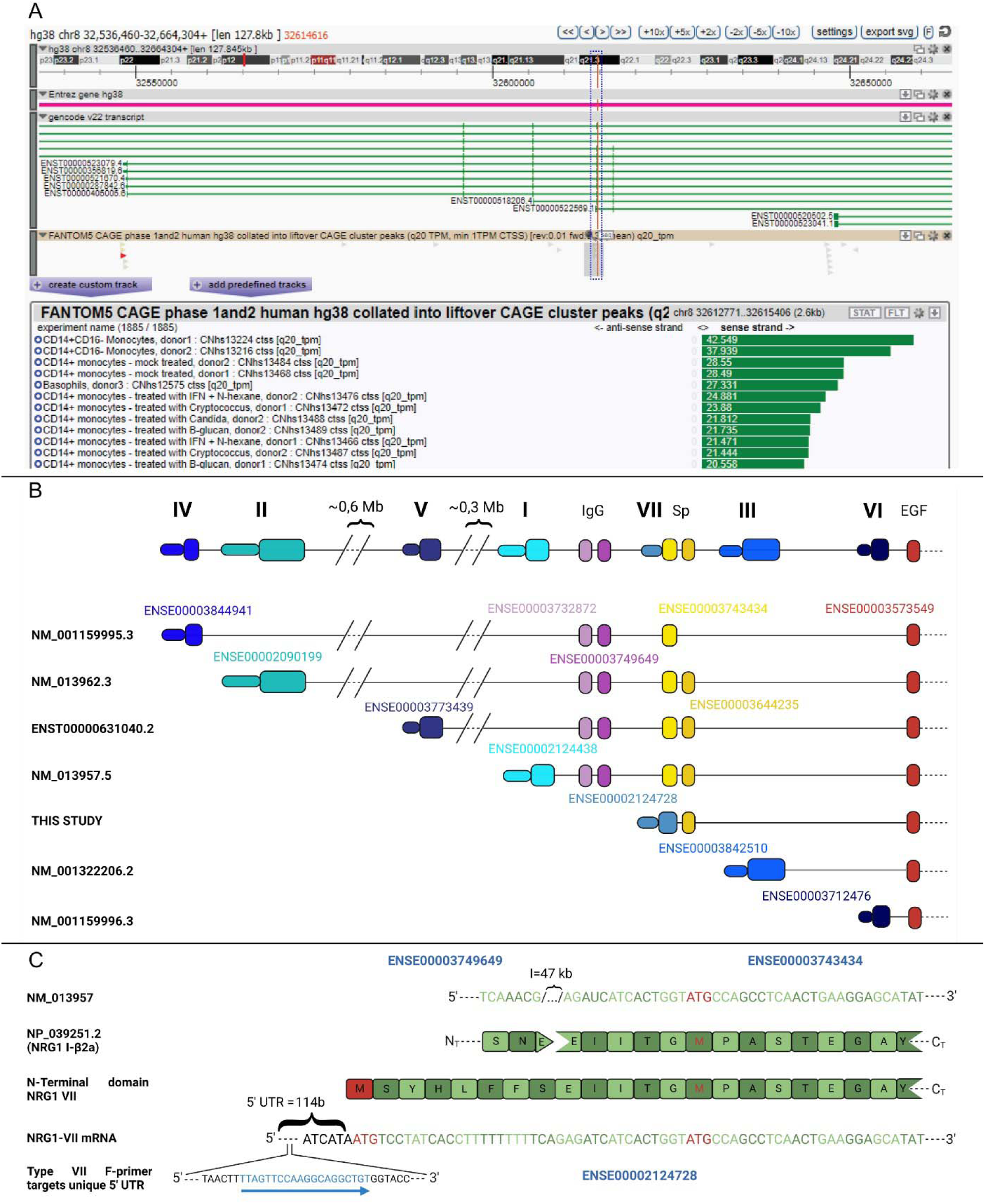
Identification of a myeloid specific TSS in the human NRG1 locus. **A)** Screenshot of the ZENBU genome browser showing the Gencode v22 transcript track, (accessed 26/1/23) corresponding to the FANTOM5 phase 1 and 2 TSS peaks overlapping the starting exon of NRG1 type VII. Red vertical line encased in blue dashed box indicates the position for the TSS of interest. Tag selection (grey bar, FANTOM5 CAGE track) indicates region for tag quantification in the FANTOM5 experiment table. The table is truncated to show the top 6 samples are primary myeloid cells, representative of the top 57/115 samples with >1TPM) Experimental samples are ordered by highest number of CAGE tags counted in this area on sense strand (green bars, showing counts as TPM). **B)** Schematic of the human *NRG1* locus highlighting alternate transcriptional starting exons that correspond to seven isoform classes. The schematic shows 5’ exon composition until the first shared exon (containing the EGF domain). Exons are numbered by ENSEMBL accessions. **C)** Alignment of mRNA and protein sequences of NRG1 type I and type VII to show an in-frame translation initiation of NRG1 type VII. Initiating methionine (M) or start codon (ATG) in red text. The blue arrow shows the target sequence of the forward primer for *NRG1-VII* mRNA’s unique 5’ UTR. The *NRG1-VII* transcript start is conserved in other mammalian genomes (Sup. Figure 1).

The TSS for NRG1-VII initiates in the intron positioned 5’ of ENSEMBL exon ENSE00003743434, which encodes for part of the spacer sequence between the Ig and EGF-like domains in the canonical NRG1 protein (Figure 2B). The TSS adds a 5’ untranslated region (UTR) to the transcript that extends the exon from 51 bp to 190 bp (ENSE00002124728) and includes an initiating Methionine at nucleotide position 115-117 bp. This newly identified TSS generates a unique 5’ UTR for its transcripts. Within this 5’-UTR, a unique 20-nucleotide sequence was identified that allowed specific detection and amplification of the NRG1-VII isoforms and primers were designed to target it (Figure 2C; Table 1). Isoforms arising from this new TSS would present a unique sequence of 8 amino acids (MSYHLFFS), a type VII specific N-terminal domain. This is followed by the 17 amino acids that are commonly present in this exon in other isoform types (I, II, IV and V). This is the shortest known isoform specific N-terminal domain, and the only one that is present within an exon that other isoforms may include.

**Table 1.**
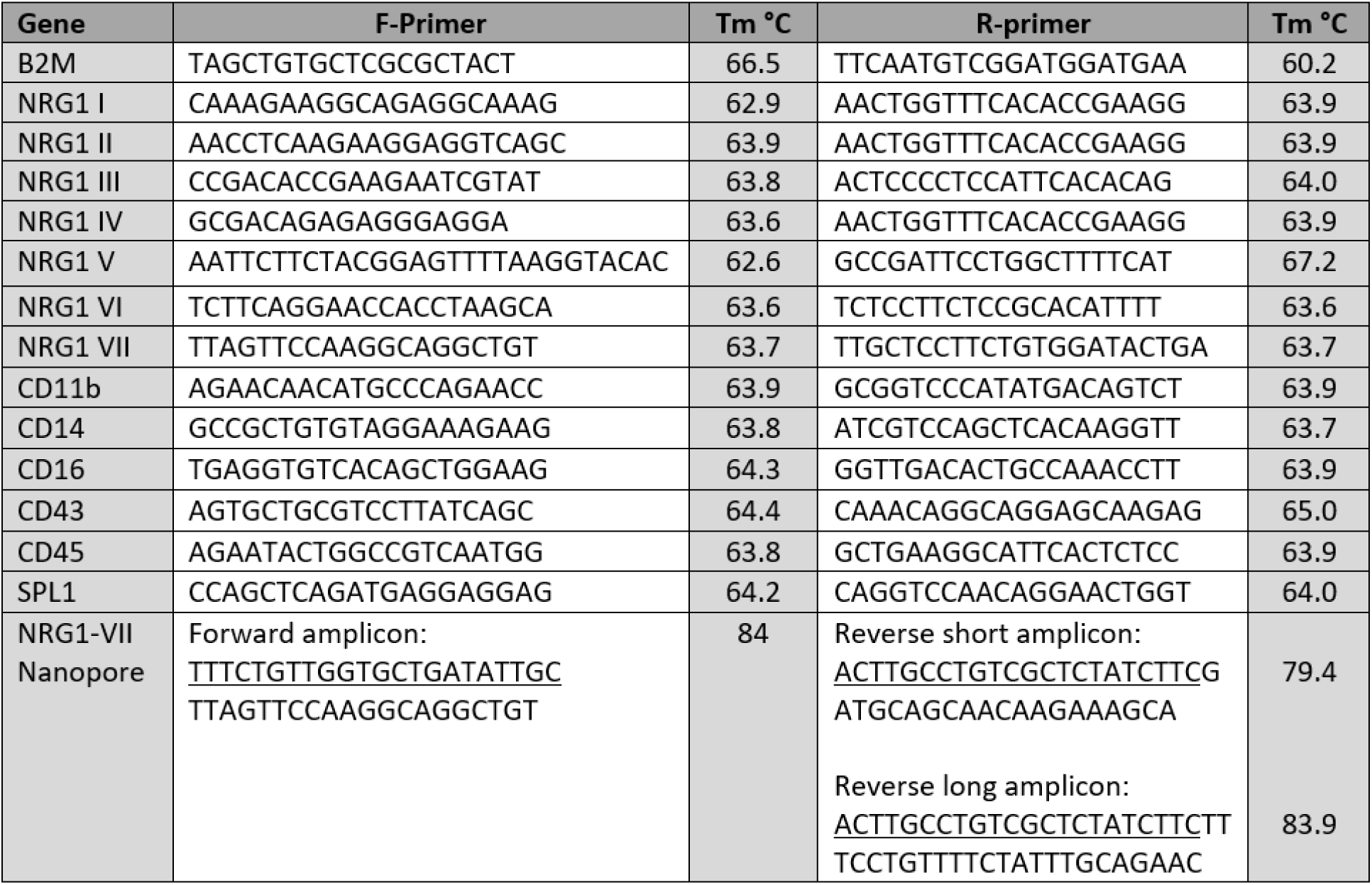
Primers designed for qRT-PCR and Nanopore Long Read Amplicon Sequencing. Note that underlined regions of the primers correspond to the general ONT 5’mod sequence.

We mapped the transcriptional activity arising from this TSS and identified 2 Expressed Sequence Tags (ESTs) that had been sequenced from the 5’ end (NCBI accessions BI908144 and BI907799) and originated from the *NRG1-VII* TSS. These originated from the same library (SAMN00164230) made from a pool of non-activated human leukocytes from anonymous donors. Both ESTs harbored evidence of an open reading frame (ORF) in frame with other NRG1 isoforms, but both were 3’ truncated. To further validate the potential activity of this TSS we looked for evidence this isoform type was present in other mammal species. Data available from 15 non-human primate species (Pipes et al., 2013), and cross species alignment shows that the *NRG1-VII* TSS is present and highly specific to bone marrow and whole blood, while isoforms isolated from other tissues use alternate TSSs (Sup. Figure 1A). We also found transcriptional evidence of an equivalent TSS in *Mus musculus* and *Sus scrofa* derived from myeloid cells (Sup. Figure 1B-C) confirming that the class VII TSS and its expression in myeloid cells are conserved through evolution.

### *NRG1-VII* TSS transcribes at least 8 distinct high-confidence transcripts in myeloid cells

To characterize the diversity of *NRG1* transcripts that use the *NRG1-VII* TSS, we performed Oxford Nanopore long-read amplicon sequencing on *in vitro* and *in vivo* derived myeloid cells (Figure 3A-E). We designed a forward primer that targets the 5’ UTR of *NRG1-VII* transcripts, which is unique to this class of NRG1 and does not overlap any other NRG1 isoform types; additionally, two reverse primers were designed to target two of the known alternative transcriptional stops that we had previously validated as active in myeloid cells through PCR (Figure 3C). We called amplicons “short” when the reverse primer targeted exon “β” and “long” for the reverse primer in exon “α”.

**Figure 3.**
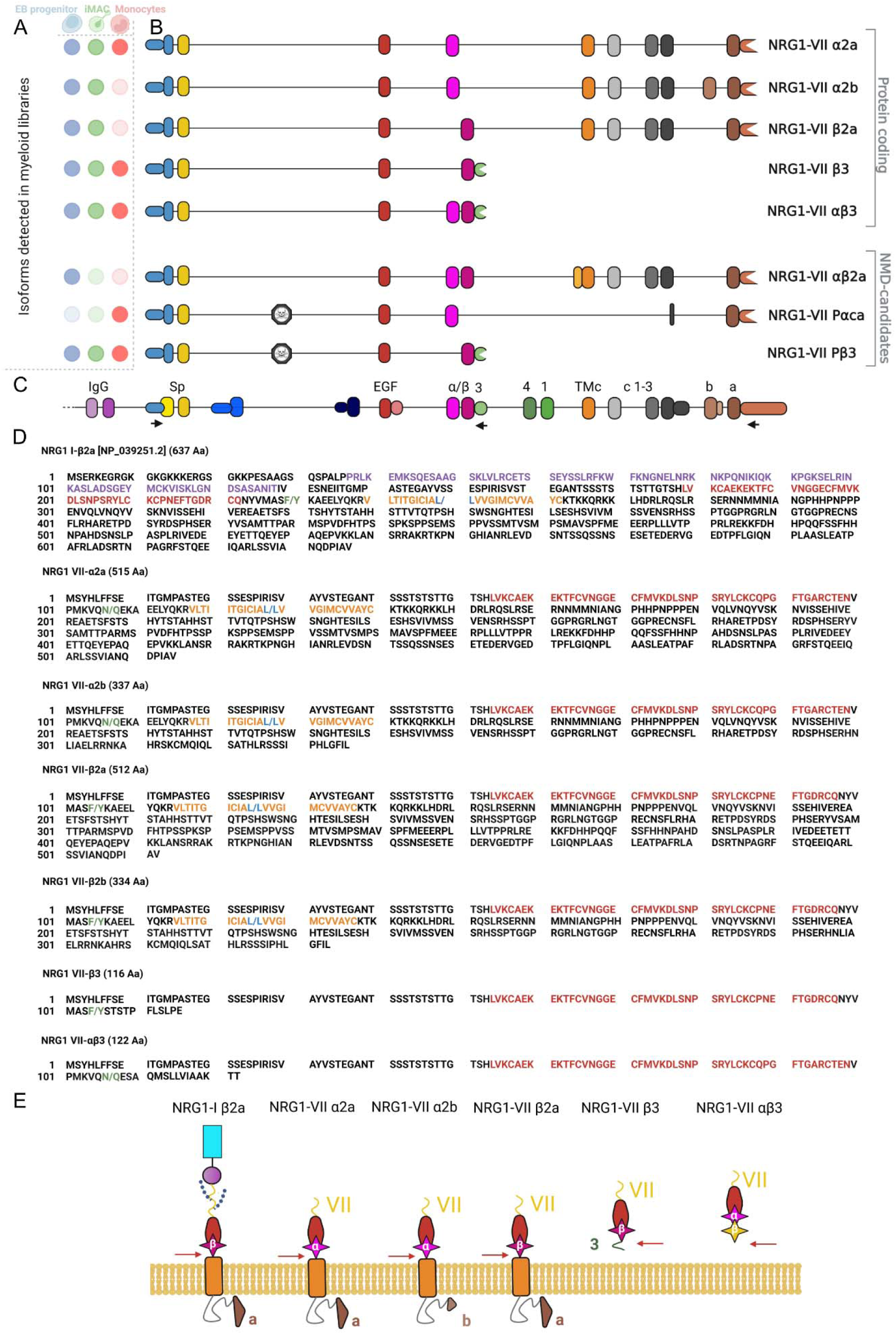
Sequence and structure of NRG1-VII isoforms. **A)** Representation of the different samples analyzed in the study and the presence or absence of the identified high-confidence transcripts in each of them (opaque = present, transparent = absent). **B)** Representation of the exon structure for 8 type VII isoforms sequenced using Oxford Nanopore long-read sequencing and their designations based on exon combinations according to the locus nomenclature. **C)** Human NRG1 locus reference with all known exons. Arrows show target regions for the single forward primer and both reverse primers used in the amplicon sequencing (See also Table 1). **D)** Predicted translation of protein sequences of the newly characterized NRG1 type VII isoforms, compared to canonical NRG1-Iβ2a (top sequence). Text colour indicates known protein motifs: purple Ig; red EGF; orange transmembrane domains. “/” represents predicted pro-peptide cleavage points (green text represents ADAM/BACE1 proteolysis, blue represents γ-secretase proteolysis). Dashed lines in all panels represent sequences that are not shown. **E)** Schematic of predicted translated proteins for NRG1-VII isoforms compared to canonical NRG1-I (Left), borrowing from domain schema shown in Figure 1. Alternate (α or β) EGF domains annotated in pink. Intracellular triangles represent alternate a or b cytoplasmic tails. The yellow β domain represents the peptide sequence that arises from the related exon but is translated to a different peptide sequence due to a frame change.

In total, we identified 8 novel high-confidence transcripts that were assigned at least 5% of the reads present in a sample from either the short or long amplicons (Figure 3B, D), using the IsoLamp pipeline (See Materials and Methods). Each amplicon was studied independently in *in vitro* progenitors and differentiated macrophages, and in blood-isolated monocytes. Five of these transcripts had an open reading frame (ORF) following a start codon (ATG), in frame with all other previously described NRG1 isoforms, and no premature stop codons in early exons. We predict that these five transcripts could be protein coding (Figure 3B), including two short and three long isoforms. The three long isoforms would include a transmembrane domain (NRG1-VII α2a, α2b and β2a) and likely undergo canonical processing through metalloproteases for the EGF-containing peptide to be released. On the other hand, the two short isoforms would lack the transmembrane and intracellular domains.

The short isoforms were detected in all myeloid libraries and lacked the transmembrane domain (Figure 3E). Of these, NRG1-VII αβ3 contained both α and β EGF-like exons, a combination that had been previously captured in refseq NM_004495, which we now show was a truncated transcript as its TSS had not been defined. This resolves the full-length sequence of the Class VII αβ isoforms. The presence of both α and β domains in NRG1-VII αβ3 introduced a frame shift in the β exon, changing the peptide sequence of this isoform to a unique C-terminal end. This feature might affect the binding dynamics of these isoforms to ERBB receptors, making it of special interest.

Monocytes expressed both short isoforms, but only expressed one long isoform, NRG-VII α2a. Two additional isoforms were detected in the monocyte library that we predict are sensitive to nonsense mediated decay (NMD). Transcripts NRG1-VII Pαca and NRG1-VII Pβ3 include a previously uncharacterized exon (Figure 3A) located in intron 2 of the other class VII transcripts. Additional support for this exon can be found in isoforms present in smaller proportions like P- α2a (Sup. Figure 2) This exon spans 111 bases (chr8:32,676,084-32,676,195). We identified this as a poison exon, as it introduces an amber stop codon (TAG), resulting in the following amino acid sequence: MSYHLFFSEIITGMPASTEGAYVSSESPIRISVSTEGANTSSFITDECCHGGQYHNTAKSICLILMF-.

A third NMD candidate was identified in the iPSC-derived myeloid progenitors, NRG1-VII αβ2a. For isoform αβ2a, this combination introduces an early stop codon that we theorize would lead the transcript to nonsense mediated decay (NMD). All three NMD-candidate transcripts passed our high-confidence transcript filters. We manually removed additional isoforms that passed our analysis threshold but had less than 5% read coverage. These isoforms varied only in a few bases across a splice junction (Sup. Figures 2-3) and were detected in only one library and were therefore considered as likely sequencing artefacts. There are two possible exceptions: NRG1-VII β2b and P- α2a. Isoform β2b was detected in both iPSC derived progenitors and iMACs, and P- α2a was found in monocytes, at levels between 1 and 5% (Sup. Figure 2).

### Macrophages are a major source of NRG1 in human tissue

We next assessed the distribution patterns of *NRG1* in single-cell RNA-seq experiments in the Human Protein Atlas (Karlsson et al., 2021), which revealed that the *NRG1* locus was actively expressed in a large variety of human tissues (Sup. Figure 4). Single cell expression data revealed NRG1 activity in different cell types, such as neurons in the brain and eye; glandular stromal cells in colon, ovary, or endometrium; endothelial cells in the heart and liver; and epithelial cells in the kidney and lung (Figure 4A). In most of these organs, macrophages are the main source of NRG1 (Figure 4A, Sup. Figure 4). One outstanding exception is the brain, where neurons show the highest levels of NRG1 expression in the human body. However, no isoform-specific single cell data is currently available to compare differential isoform expression between the different macrophage types present across the tissues investigated.

**Figure 4.**
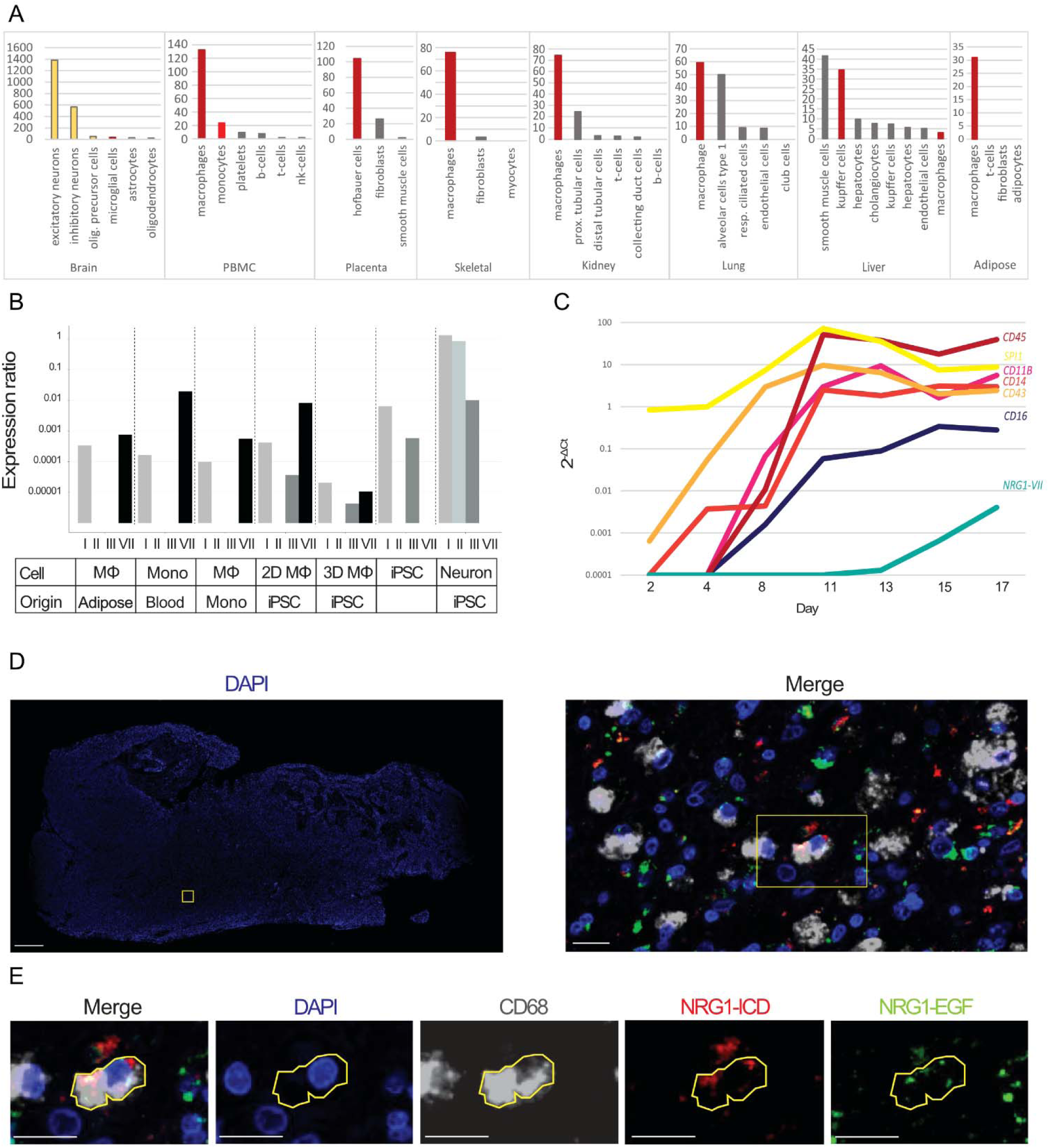
NRG1 expression profile in myeloid cells. **A)** Quantification of NRG1 mRNA expression in different human tissues according to The Human Protein Atlas (Karlsson et al., 2021). A plethora of representative tissues that express NRG1 are shown here. The y axis represents nTPM (Number of transcripts per million). Neurons shown in yellow, myeloid cells in red, other cell types in grey. **B)** qRT-PCR data measuring presence of the different NRG1 classes in different cell types. Classes IV-VI were not detected in any of the samples. Expression ratios were calculated as described in Pfaffl, 2021. Mono= Monocyte, Mφ: macrophage, iPSC: Induced pluripotent stem cell. **C)** Time series showing qRT-PCR expression of typical myeloid markers in progenitor maturation and relationship with myeloid specific NRG1-VII transcripts. **D)** Immunostaining of GBM showing presence of CD68^+^ cells that express NRG1 peptides *in vivo*. The left panel shows sample section stained with DAPI. The boxes indicate the regions shown in the right panel in D and the cell in E, respectively. Scale bar on left panel is 1 mm, scale bar on right panel represents 20 μm. **E)** Example of cell in which all markers (CD68, NRG1-EGF and NRG1-ICD) are expressed. Scale bars = 20 μm.

### NRG1-VII expression in myeloid cells is affected by differentiation and maturation

We sought to confirm if isoform VII is the only or even primary TSS used by myeloid cells. Therefore, using primers that could discriminate between each unique start exon (Table 1), we investigated the patterns of the seven classes of NRG1 isoforms in different *in vivo* and *in vitro* myeloid cells (Figure 4B). Expression of *NRG1-VII* was detected in all myeloid cells, and it was the isoform class showing highest expression for cells that belong to this lineage (except for macrophages derived from a 2D epithelium). In contrast, control cell types (hiPSCs and iPSC-derived cortical neurons) showed high expression of other *NRG1* classes, but not *NRG1-VII*.

Having demonstrated that our iPSC derived myeloid cells can be used to model the use of this TSS, we next investigated at which differentiation stage we could first detect the expression of NRG1-VII transcripts. Thus, we investigated the NRG1-VII expression patterns during the differentiation process towards the myeloid lineage, and its concurrence with specific myeloid markers (Figure 4C). Our results show that NRG1-VII transcription in *in vitro* derived myeloid cells follows the upregulation of mature monocyte markers like CD16, unlike what is observed in blood monocytes from *in vivo* samples (Figure 4C). This suggests that the sequence of events happening *in vivo* are not recapitulated in our *in vitro* differentiation, and that a more mature myeloid identity is needed to activate transcription arising from the *NRG1-VII* TSS in these cells.

### NRG1-VII expression in myeloid cells is affected by differentiation and maturation

To confirm that *NRG1* mRNAs are translated into proteins in myeloid cells, we then performed immunohistochemical staining of NRG1 in human glioblastoma (GBM) tissue which is enriched in bone marrow-derived macrophages (Klemm et al., 2020). Myeloid cells were identified using a CD68 antibody; antibodies detecting the extracellular (EGF-like domain) and intracellular (ICD) domains of NRG1 were used. NRG1 was detected in both myeloid and non-myeloid cells (Figure 4D-E). Not all CD68+ cells expressed NRG1, and those cells that did exhibited different patterns of expression (Sup. Figure 2). Altogether, these results show that myeloid cells express NRG1 peptides in human tissues but are not the sole contributors to the NRG1 pool in the brain. They also reveal diverse NRG1 expression patterns in myeloid cells, which may be consistent with the different isoforms observed in our sequencing libraries, or consistent with active processing of membrane bound-NRG1.

## Discussion

Macrophages have been reported as a major source of NRG1 in different human tissues (Karlsson et al., 2021), including a recent report highlighting the importance of NRG1 in the developing gut (Jardé et al., 2020). Our investigation of *NRG1* expression in the public single cell atlases, including the Human Protein Atlas mRNA dataset, suggested that macrophages are a major source of NRG1 in most tissues. However, the specific isoforms involved in the macrophage mediated NRG1 secretion were uncharacterized. Here we report that myeloid cells preferentially use a novel TSS that is myeloid-specific and conserved which generates a previously uncharacterized class of NRG1 isoforms. The tissue variability and functional versatility of NRG1 are a consequence of the alternative promoter usage and exon retention that give rise to a high diversity of isoforms.

Transcripts arising from the human *NRG1* locus are differentially regulated by tissue type or developmental stage. Control over the six known isoform classes is achieved through differential proximal promoter usage at six unique TSSs, each of which is controlled by different transcription factors (Frensing et al., 2008). All available data indicates that the use of this TSS is exclusive to the myeloid lineage. The concrete mechanisms and transcription machinery involved in the process are yet to be described. Additional transcript processing can lead to alternative exon usage and define the functional modules in the protein, like the linker or the cytoplasmic tails (Figure 1 C, D), mechanisms that are tissue-specific (Wen et al., 1994). The diversity of NRG1 isoform classes and their specificity of expression in both a temporal anatomical and cell-specific manner, suggest distinct roles of different isoforms in tissue patterning, especially in different brain regions (Liu et al., 2011). Further characterization of each of the NRG1-VII isoforms is hence needed to elucidate their regulation and functional diversity.

It was previously reported that the highest NRG1 expression levels in human tissues was in blood plasma (Pipes et al., 2013). This is unsurprising, as maturation of myeloid cells *in vivo* seems to cause the downregulation of this gene across the monocyte-macrophage differentiation axis (Figure 2A), with a subset of macrophages expressing NRG1 in tissues (Figure 4). Our review of the FANTOM and The Human Protein Atlas suggests that monocytes are the main source of NRG1 in blood. The function of monocyte-derived NRG1 has not yet been described, however, circulating levels of NRG1 correlate with liver metabolic activity (Zhang et al., 2018) as well as post-infarct recovery and cardiovascular health improvement (Mendes-Ferreira et al., 2013). This suggests that circulating NRG1 could play an important role in hepatic and cardiac health.

We further found that monocytes express at least five high-confidence NRG1-VII isoforms, including transcripts that contain a novel ‘poison’ exon (NRG1-VII P αca and Pβ3) predicted to prematurely terminate translation (Figure 3A). We also note the unusual exon composition of isoform NRG1-VII Pαca, indicating it could be a PCR artifact; thus, further validation might be required on this isoform, despite its amplicon constituting over 15% of the detected expression. Additional isoforms containing the monocyte poison exon like NRG1-VII Pα2a were also found, but in much lower proportions (Sup. Figure 2). However, this supports the use of the poison exon by monocytes. This exon introduces an early stop codon and hence is likely to drive the transcript to nonsense mediated decay, generating a monocyte specific transcriptional mechanism to regulate the levels of NRG1 synthesis. Due to the high levels of expression of this gene in monocytes compared to other cell types, we hypothesize that these represent regulatory mechanisms that allow control over the levels of NRG1 expression in circulation.

Neuregulins interact with monomeric ERBB proteins (ERBB1-4), via an EGF-like domain, promoting dimerization and trans-phosphorylation of these receptors. Each ERBB monomer can bind different EGF members with differential ligand binding affinities, but only ERBB 3 and 4 can bind Neuregulins. Though NRG1 and NRG2 can bind to both receptors as monomers (Carraway III et al., 1997), NRG3 and NRG4 only bind ERBB4 monomers (Harari et al., 1999, Zhang et al., 1997). Dimerized receptors can discriminate between different isoforms of NRG1, which is evidenced by differential phosphorylation on receptor tyrosine residues (Pinkas-Kramarski et al., 1998, Sweeney et al., 2000). Dimerization leads to the recruitment of different adaptor proteins, notably GRB or PI3K, which then trigger specific secondary cascades and regulate cell activity (Buonanno et al., 2001). NRG1-VII isoforms present a very short N-terminal domain compared to most other isoforms. It has been previously reported that the size and structure of the NRG1 N-terminal domain can dictate the receptor availability and recycling, affecting internal phosphorylation and internal signaling in the receiving cell (Warrant et al., 2006). Additionally, the discovery of isoform NRG1-VII αβ3 as a coding isoform, adds a novel sequence possibility in the EGF domain. This is likely to determine the protein’s binding dynamics and internal phosphorylation, hence changing its functional properties. Functional validation is necessary to confirm whether this isoform type has unique properties given its EGF domain structure.

NRG1 is involved in many different functions, attributed to the many isoforms which are tissue-restricted during different developmental stages. Further, while NRG1 is conserved at the protein level, the function of specific isoforms has been shown to differ between species. For example, heart trabeculation failure, which leads to a lethal phenotype in mice at day 10.5, is a phenotype common to NRG1-/-, ERBB2-/- and ERBB4-/- mice (Kramer et al., 1996, Britsch et al., 1998). Although 11 different isoforms have been identified in the heart at this developmental stage, the phenotype can be attributed specifically to the absence of β-RGF types I and II, which are the isoforms containing the immunoglobulin (Ig) domain (Pentassuglia et al., 2009). However, the opposite is observed in zebrafish where Ig-like neuregulin-1 isoforms are dispensable for this process (Samsa et al., 2016). Thus, while essential processes in development can be attributed to specific subtypes of NRG1, differences between species showcase the need to characterize the different human isoforms using appropriate cell type and developmental models.

NRG1 and the ERBB receptors are important clinical target in different cancer types. For example, NRG1 fusion proteins cause pathological activation of ERBB receptors (Laskin et al., 2020). NRG1 expression correlates with a shorter survival in patients with glioma (Yin et al., 2015), which could be related to its role in cell migration and proliferation. Glioblastomas particularly exhibit high macrophage infiltration (Wei et al., 2019). Our results show that tumor-associated myeloid cells (CD68^+^) may contribute to the NRG1 pool seen in these tumors. Due to the lack of unique domains in type VII isoforms, the antibodies used in this study targeted common NRG1 regions (the EGF-like and intracellular domains). Even though the available antibodies lack specificity to prove that type VII isoforms are translated in this disease, the only isoforms detected in primary monocytes or macrophages in this study belong to classes I and VII.

While NRG1 double positive myeloid cells may express newly synthesized pro-peptides, we also observed CD68^+^/NRG1-ICD^+^ cells indicating that the EGF-like domain had been cleaved. Alternatively, the presence of CD68^+^/NRG1-EGF^+^ cells suggests that some cells restrict their expression only to isoforms that end transcription in linker 3 and lack a transmembrane domain. Therefore, we confirmed that macrophages *in vivo* express NRG1 and that there are distinct populations of macrophages in tissue based on their NRG1 expression patterns.

To determine whether *in vitro* derived myeloid cells can be used to model the activity of the NRG1-VII TSS, we used primers designed to target specific TSS usage in different cell types. We confirmed that NRG1-VII was expressed in all myeloid samples and in no others, suggesting the TSS is active. However, iPSC-derived samples showed expression of NRG1-III. This could be a result of incomplete differentiation of the culture, or retention of stem cell features in *in vitro* derived myeloid cells. Moreover, the time series data obtained during the differentiation process shows that NRG1 expression follows mature markers like CD16; *in vivo*, NRG1 expression precedes CD16, and as CD16 is upregulated, NRG1 expression decreases. Hence, while we show activity of the TSS, the *in vitro* model may not recapitulate the transcriptional and maturation sequence seen *in vivo* due to the differences in the differentiation process.

Description of a new TSS class, NRG1-VII, including at least five new protein-coding isoforms, has expanded the known NRG1 protein coding isoforms from 28 to 33. This study adds eight new transcripts that are specific to myeloid cells. However, it is likely that the full transcriptional profile of this locus has not yet been described; for example, while long-read amplicon sequencing is a sensitive isoform recovery method (Clark et al., 2020), it is limited by the primer set(s) used. Thus, only NRG1-VII isoforms utilizing linker 3 or the “a tail” regions were amplifiable in this study. It is highly likely that NRG1-VII isoforms using the 3’ end present in exon c3 (ENSE00002109887) also exist, which should be validated with additional work.

Here we showed that myeloid cells exhibit a unique regulation pattern of the NRG1 locus to generate cell specific isoforms, potentially playing an important role in diverse diseases. Only through a thorough investigation of this locus can we better understand each process and develop clinical strategies to prevent or treat the different pathologies associated with this locus. Further detailed investigation on the molecular genetic features and functions of these novel isoforms might uncover how NRG1-VII isoforms elicit differential receptor activity and downstream effects as previously described for other NRG1 isoforms. This could lead to targeted therapies and an improved understanding of the complexity of the *in vivo* system, helping us recreate the processes in which appropriate signals are essential to model the desired biological mechanisms.

## Materials and methods

### Cell lines and ethics approvals

Stem cell work was performed in accordance with The University of Melbourne ethics committee HREC (approval 1851831). The line of human iPSCs used was: PB001.1 (Vlahos et al., 2019), obtained from the Stem Cell Core Facility at the Murdoch Children’s Research Institute. Kolf2.1 (hPSCReg accession WTSIi018-B; Welcome Trust Sanger Institute) cells were used to differentiate cortical neurons (Ethics ID: 12374) and 2D macrophages. Monocytes were isolated from buffy coat, which was obtained from the Australian Red Cross Blood Service in accordance with The University of Melbourne ethics committee HREC (approval 1646608). Ethics for adipose tissue derived samples was obtained from the University of Melbourne Human Ethics Committee (ethics ID 1851533) and approved by The Avenue Hospital Human Research Ethics Committee (Ramsay Health; ethics ID WD00006, HREC reference number 2019/ETH/0050). For human glioblastoma samples, human ethics approval was covered by project application 1853511, approved by the Medicine and Dentistry Human Ethics Sub-Committee, The University of Melbourne.

### Stem cell culture

Cells were cultured in Gibco™ Essential 8™ media with Essential 8™ supplement (Thermo Fisher Scientific; A1517001) on growth-factor reduced Matrigel® Matrix (Corning®; 356234) coated dishes. Cells were cultured with daily changed fresh media. Cell culture was performed in an APT.line^TM^ C150 (E2) CO_2_ manual incubator (BINDER; 7001-0172) in constant and stable conditions of humidity, temperature (37°C) and CO_2_ concentration in air (5%).

Cell passaging was performed routinely when cell confluency reached (70-80%). Cells were first washed with Gibco™ PBS (Thermo Fisher Scientific; 10010023). Then, a dilution of sterile 0.5 M EDTA (Thermo Fisher Scientific; 15575020) in said PBS at a concentration of 0.5 mM was used to detach the cells from the plate. After 3-4 minutes in humid incubator conditions, cells were collected and replated with fresh media.

### 3D iPSC derived macrophages

As described in Rajab et al., 2021. Human iPSCs were differentiated into macrophages following (Joshi et al., 2019), but with the following alterations to the protocol: harvested cells were cultured in MAGIC media, which was changed as indicated in (Ng et al., 2008). During this process, cells were plated in 10cm Ø non-treated Petri dishes (IWAKI; 1020-100) and placed on an orbital shaker (N-Biotek orbital shaker NB-T101SRC) in a humidified incubator with 5% CO_2_ at 37°C.

After 11 days of culture, cells can be observed detaching from the embryoid bodies, remaining in suspension in the media as non-adherent cells (which are characterized as myeloid progenitors). When all cells are collected and allowed to settle in a 15 mL Falcon tube (Corning®; CLS431470-500EA), embryoid bodies pelleted in the bottom, but progenitors stayed in suspension in the supernatant, and could then be collected. The supernatant was then centrifuged (Heraeus Multifuge 1S-R) at 400 rpm for 5 minutes to separate the progenitors from the media. Progenitors were then resuspended in a 10% dilution in volume of FBS and 100ng/mL CSF-1 (R&D Systems; 216-MC-500) in Gibco™ RPMI-1640 media (Thermo Fisher Scientific; 11875093). Cells were plated in Costar® 6-well tissue-culture treated plates (Corning®; 3516) for 4-7 days in stable incubator conditions (humid, 5% CO_2_, 37°C), when cells showed morphological and molecular features displayed by macrophages.

### iPSC derived cortical neurons

Kolf2.1 hiPSCs (Kao et al., 2016) were cultured under xenogenic conditions as defined in Niclis et al. (2017). The cells were then differentiated into cortical neurons following the protocol described by (Gantner et al., 2021).

### Human samples and cell sorting

#### Blood monocyte isolation

Buffy Coat was obtained from the Australian Red Cross Blood Service. The blood was diluted with PBS at a 1:3 dilution and underlaid with Histopaque®-1077 (Sigma-Aldrich; Cat. No. 10771-100ml). The underlaid blood samples were centrifuged (TECHCOMP CT1SRT) at 350g for 30 minutes at 24°C with no brake. Peripheral blood mononuclear cells (PBMCs) were isolated from the interphase and washed twice using MACs buffer (Gibco™ Dulbecco’s phosphate-buffered saline (DPBS) (Ca2+Mg2+ free) (Thermo Fisher Scientific; Cat. No. 14190144) with 0.5% heat inactivated Fetal Bovine Serum (FBS) (Thermo Fisher Scientific; Cat. No. 10082147 or 10099141) and 2mM EDTA (Invitrogen™ UltraPure™ 0.5M EDTA (Thermo Fisher Scientific; Cat. No. 15575020)) and centrifuging at 400g for 5 minutes at 4°C. Cell count and viability were determined using 0.4% Gibco™ Trypan Blue (Thermo Fisher Scientific; Cat. No. 15250061) using a hemocytometer. Cells were centrifuged at 400g for 5 minutes at 4°C and resuspended in 40μl MACs buffer per 10^7^ cells. Monocytes were positively selected by a magnetic field using Human CD14 MicroBeads (Miltenyi Biotec; Cat. No. 130-050-201) and LS Columns (Miltenyi Biotec; Cat. No. 130-042-401).

#### Adipose tissue samples

Adipose tissue was obtained and processed as described in (Raajendiran et al., 2019). Myeloid cells were selected by FACS using a BD FACSAria™ III system (BD Biosciences). Cells that were positive for markers CD45 and CD11b were isolated for this study.

### RNA extractions and cDNA synthesis

Total RNA was extracted using the RNeasyⓇ Plus Mini Kit (Qiagen; 74134) according to manufacturer’s instructions. After final total RNA elution in RNase-free water, overall RNA quality and concentration were measured using an RNA ScreenTape (Agilent Technologies; 5067-5576) in an Agilent 2200 Tapestation System (Agilent Technologies; G2964-90003).

cDNA synthesis was then performed using the isolated total RNA and considering the concentration values assigned to each of the samples for the coming steps. cDNA was synthesized using SensiFASTTM cDNA Synthesis Kit (BioLine; BIO-65053) and following all protocol specifications from the vendor. cDNA concentration and quality were checked using a D5000 ScreenTape (Agilent Technologies; 5067-5588) in an Agilent 2200 Tapestation System. Final cDNA samples were then stored at −20°C.

### q-RT PCR

For mRNA quantification, the Applied Biosystems Viia7 ™ real time system was used (Thermo Fisher Scientific; 4453536) using a Fast 96 well plate hardware set up. The reactions in each well of the Micro Amp Fast 96 well reaction plate 0.1 mL (Thermo Fisher Scientific; 4346907) were prepared as indicated for the Fast SYBR™ Green Master Mix (Thermo Fisher Scientific; 4385612). Primers used in the reaction were designed for each transcript class of interest (Table 1). Plates were then sealed using Optical adhesive covers (Thermo Fisher Scientific; 436 0954). For quantification, n=3 technical replicates were used.

Relative expression ratio was calculated as E(HKG)Average HKG Ct/ E(GOI)Average GOI Ct a described in Pfaffle, 2021. Efficiencies for each primer pair were calculated from serial dilutions of template and ranged between 95 and 98 % for all NRG1 isoforms classes reported in results. Isoforms IV, V, VI were not detected in any samples. B2M was used as House Keeping gene (HKG) for all samples.

### Nanopore Amplicon sequencing

NRG1-VII was amplified using the LongAmp® Taq 2X Master Mix (New England Biolabs; Cat #: M0287S) and the specified primers (Table 1) for either 30 cycled for the shorter amplicon or 40 cycles for the longer amplicon. The samples used were a pool of monocytes from 3 different blood donors, a pool of myeloid progenitors derived *in vitro* and macrophages differentiated from said *in vitro* derived progenitors. Target sequences were then amplified using specific primers (Table 1) and purified using AMPure beads (Beckman Coulter; A63880) at concentrations appropriate to each of the target sizes. Then, 2 ng of purified cDNA from each sample were barcoded following the EXP-PBC096 protocol from Oxford Nanopore Technologies. Samples were pooled (equimolar) and a sequencing library was prepared as described in the SQK-LSK110 Oxford Nanopore protocol. Samples were then loaded on a Flongle flow cell (FLO-FLG001) and sequenced using a GridION device.

### Sequencing data analysis

Basecalling was performed using Guppy (v6.3.8) using the super-accurate basecalling config file with a Qscore threshold of 10. To identify NRG1 Isoforms we used IsoLamp v1.0 (https://github.com/ClarkLaboratory/IsoLamp), a bash pipeline for the identification of known and novel isoforms from targeted amplicon long-read sequencing data generated with Oxford Nanopore technologies. Briefly, IsoLamp takes passed Nanopore reads and maps them to a reference genome using Minimap2 (Li, 2018). Alignments that are highly accurate (>95%), are full-length and have high accuracy splice junctions (>90%) are used for isoform discovery with bambu (v3.2.4). Isoform quantification is then performed with Salmon v0.14.2 (ref). IsoLamp was run in de novo mode setting BambuAnnotations=NULL. Each sample was run independently through the IsoLamp pipeline with ‘downsample reads’ parameter set to FALSE. All other parameters were set to default. Isoform annotation files and count matrixes were visualized using IsoVis (https://isomix.org/isovis).

### Automated multiplex immunohistochemistry

Formalin-fixed paraffin-embedded (FFPE) GBM tissue sections were stained using the Bond RX automated stainer (Leica Biosystems). Slides were deparaffinized in xylene followed by exposure to a graded series of ethanol solutions for rehydration. Heat-induced epitope retrieval was performed with either a Citrate pH 6 buffer or Tris Ethylenediaminietetraacetic acid (EDTA) pH 9 buffer. Slides were blocked with 3% hydrogen peroxide (H_2_O_2_) to block endogenous peroxidase activity. For multiplexed IHC staining the Opal 6-plex Detection Kit (Akoya Biosciences) was used. Serial multiplexing was performed by repeating the sequence of antigen retrieval, primary antibody, and Opal polymer incubation, followed by Opal fluorophore visualisation for all antibodies as follows. GBM tissue was stained with CD68 (Abcam ab955, 1:100), NRG1 ICD (Abcam ab191139, 1:200) and NRG1 EGF-like domain (ThermoFisher; MA4-12896, 1:100). Slides were incubated for 1 hour at room temperature with primary antibodies diluted in 1x Opal blocking/antibody diluent (Akoya biosciences). Slides were subsequently incubated with the Opal Polymer HRP Ms + Rb secondary polymer for 30 minutes prior to incubation with Opal fluorophores (Opal 520, 540, 570, 620, 650 and 690) diluted at 1:150 in 1x Plus Automation Amplification Diluent (Akoya biosciences) for 10 minutes. Slides were counterstained with 10x Spectral DAPI and coverslipped with ProLong Glass Antifade Mountant (Invitrogen). Multispectral images were acquired at 20x and 40 magnification using PhenoImager ™ HT (Akoya Biosciences). inForm 2.4.8 software (Akoya Biosciences) was used for spectral deconvolution. Deconvoluted multispectral images were subsequently fused in HALO (Indica Labs).

## Supplementary data

**Supplementary Figure 1.**
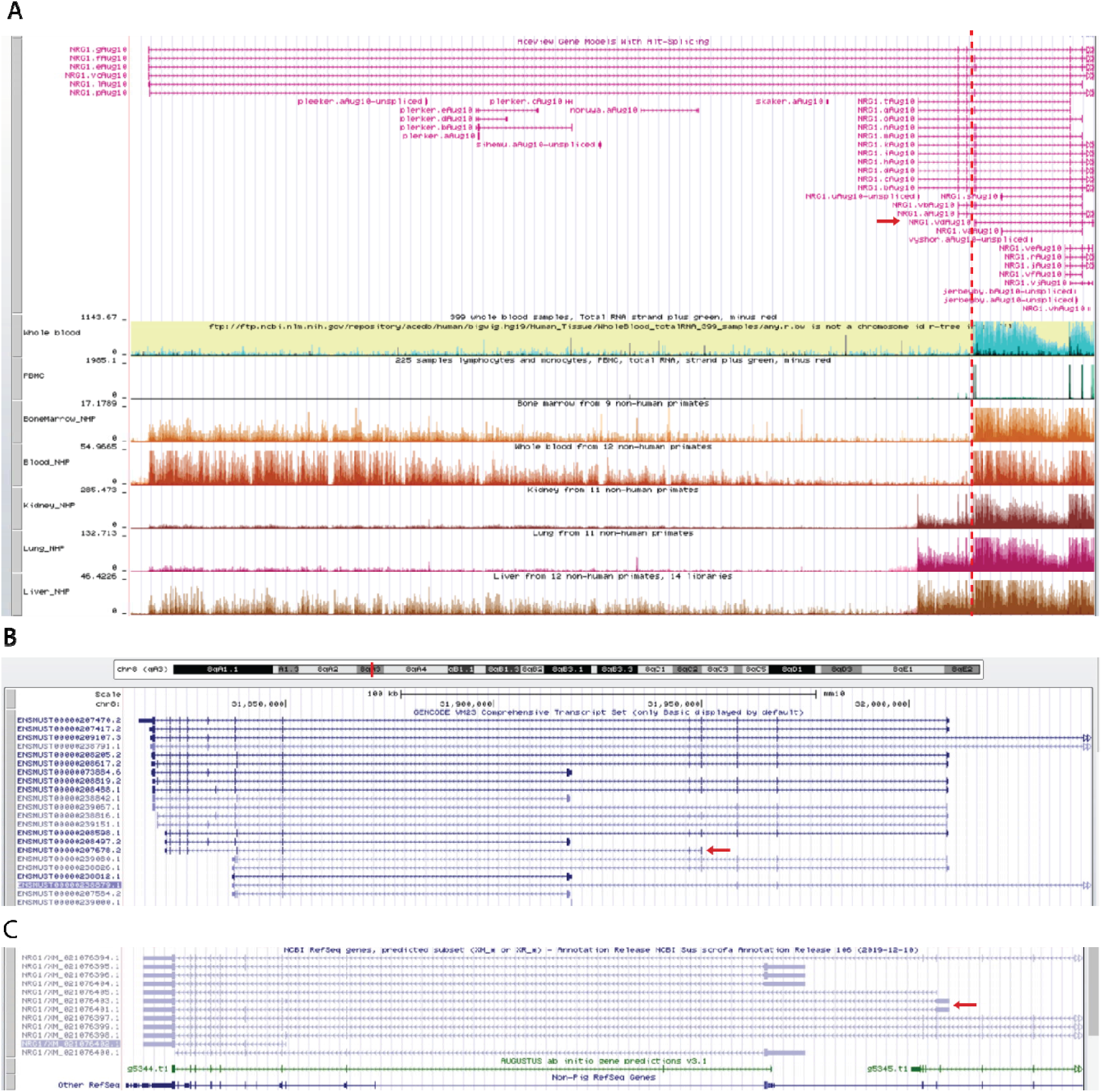
NRG1-VII TSS shows expression conservation in primate and non-primate mammals. **A)** NRG1 expression profiles of human and non-human primates showing that TSS VII is conserved in bone marrow and whole blood **B)** Curated transcript in *Mus musculus* database showing conservation of TSS VII in blood cells. **C)** Predicted transcript in *Sus scrofa* based ETSs derived from dendritic cells and other myeloid progenitor samples. Screenshot from https://www.ncbi.nlm.nih.gov1. *Accessed on 15/12/2022*.

**Supplementary Figure 2.**
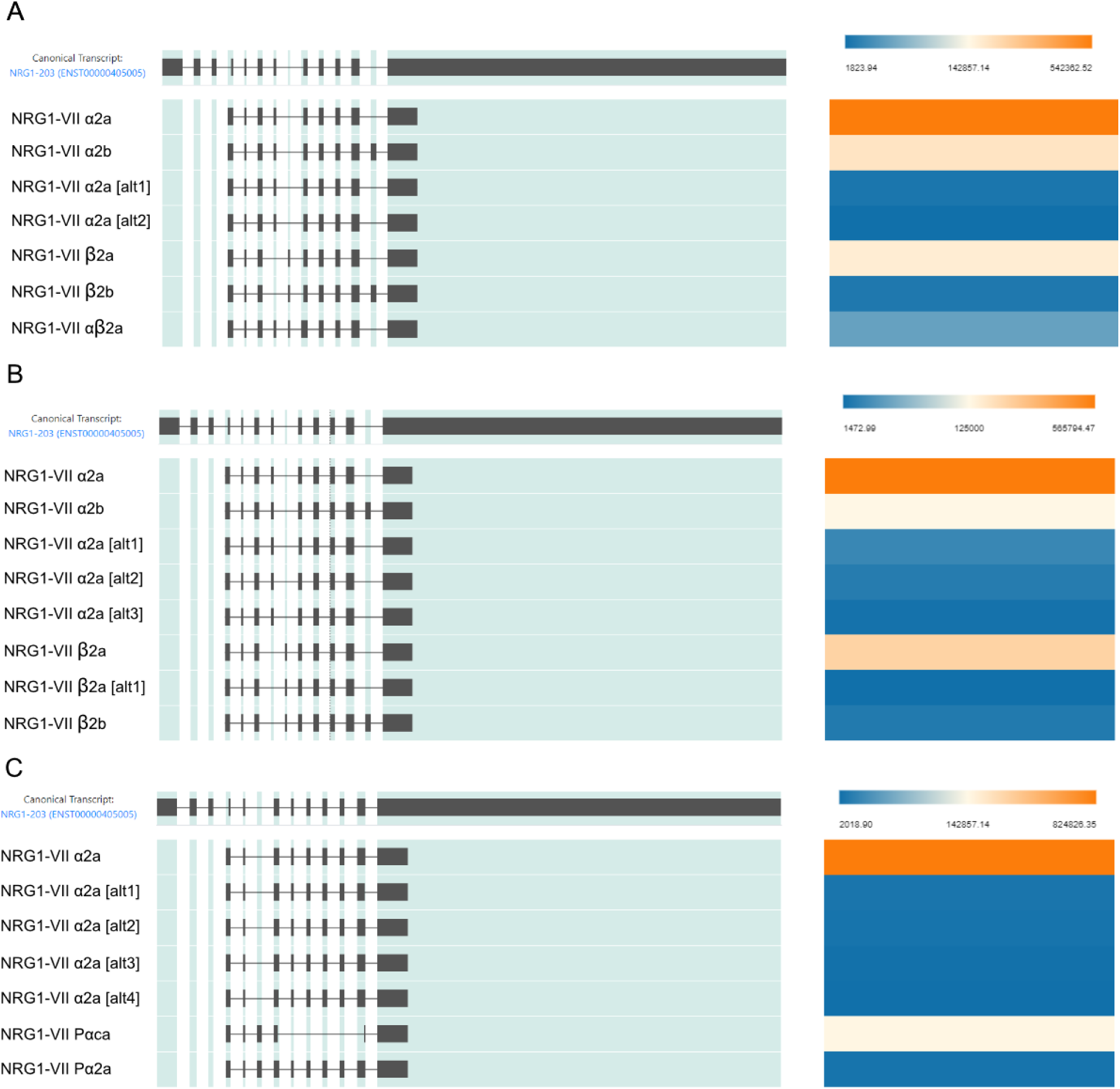
NRG1 long amplicon isoforms found through IsoLamp. Each panel shows the IsoVis (Ref) view for NRG1 amplicons in a different sample. The initial row shows the canonical isoform structure chosen by Isomix. The rest show the resulting isoforms found by the IsoLamp analysis pipeline. Isoforms that include the term “[alt]” show isoforms that are only different to the main isoform in one of the splicing junctions by a few bases. The heat maps on the right-hand side describe relative abundance of each transcript in that sample in nTPM. Panels correspond to **A)** iPSC derived myeloid progenitors; **B)** iPSC derived macrophages (iMACs); **C)** Monocytes.

**Supplementary Figure 3.**
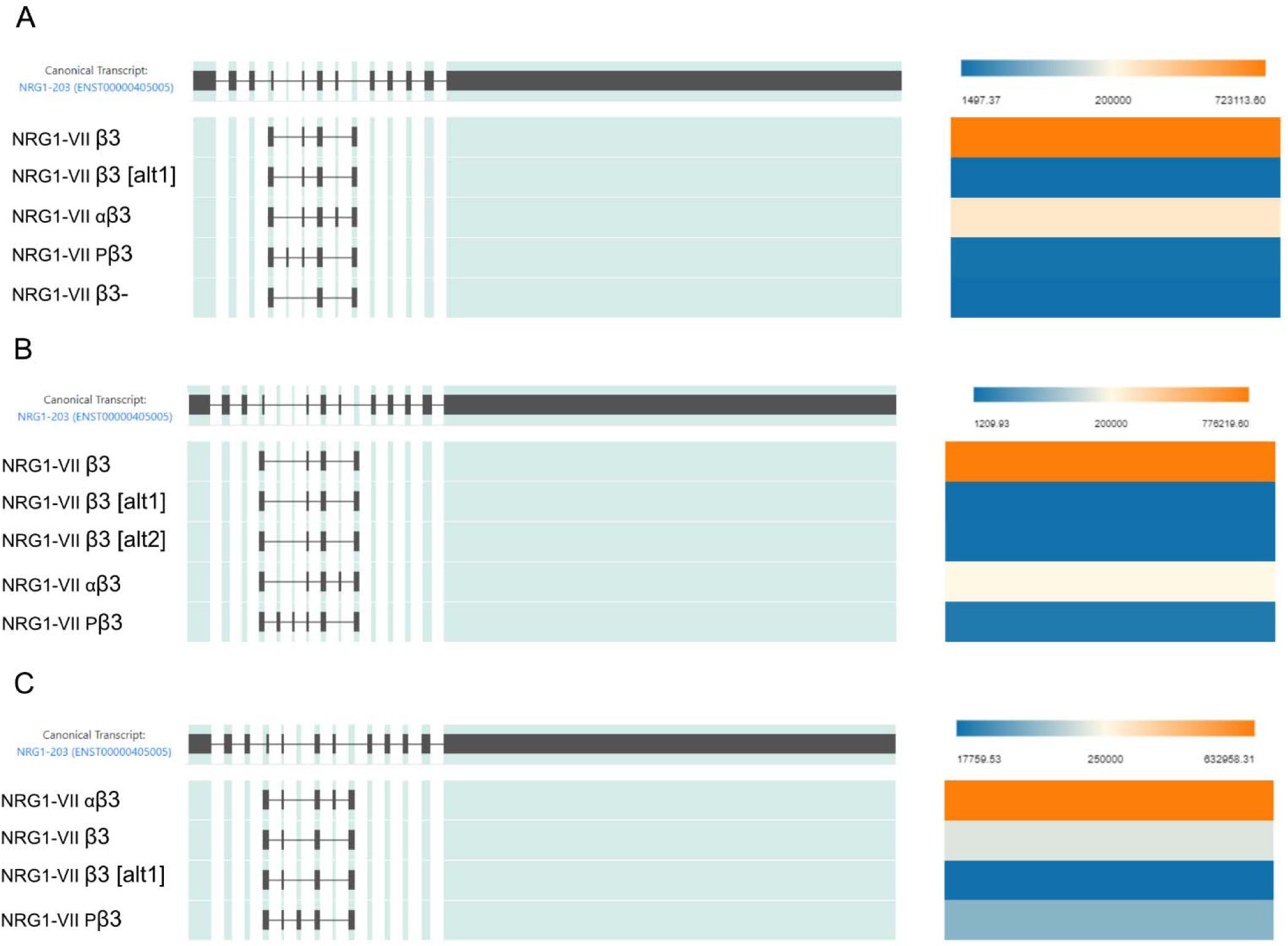
NRG1 short amplicon isoforms found through IsoLamp. Each panel shows the IsoVis (Ref) view for NRG1 amplicons in a different sample. The initial row shows the canonical isoform structure chosen by Isomix. The rest show the resulting isoforms found by the IsoLamp analysis pipeline. Isoforms that include the term “[alt]” show isoforms that are only different to the main isoform in one of the splicing junctions by a few bases. The heat maps on the right-hand side describe relative abundance of each transcript in that sample in nTPM. Panels correspond to **A)** iPSC derived myeloid progenitors; **B)** iPSC derived macrophages (iMACs); **C)** Monocytes. Different poison exons were detected for isoform NRG1-VII Pβ3 in each sample type, therefore further validation of these exons and isoform(s) is needed.

**Supplementary Figure 4.**
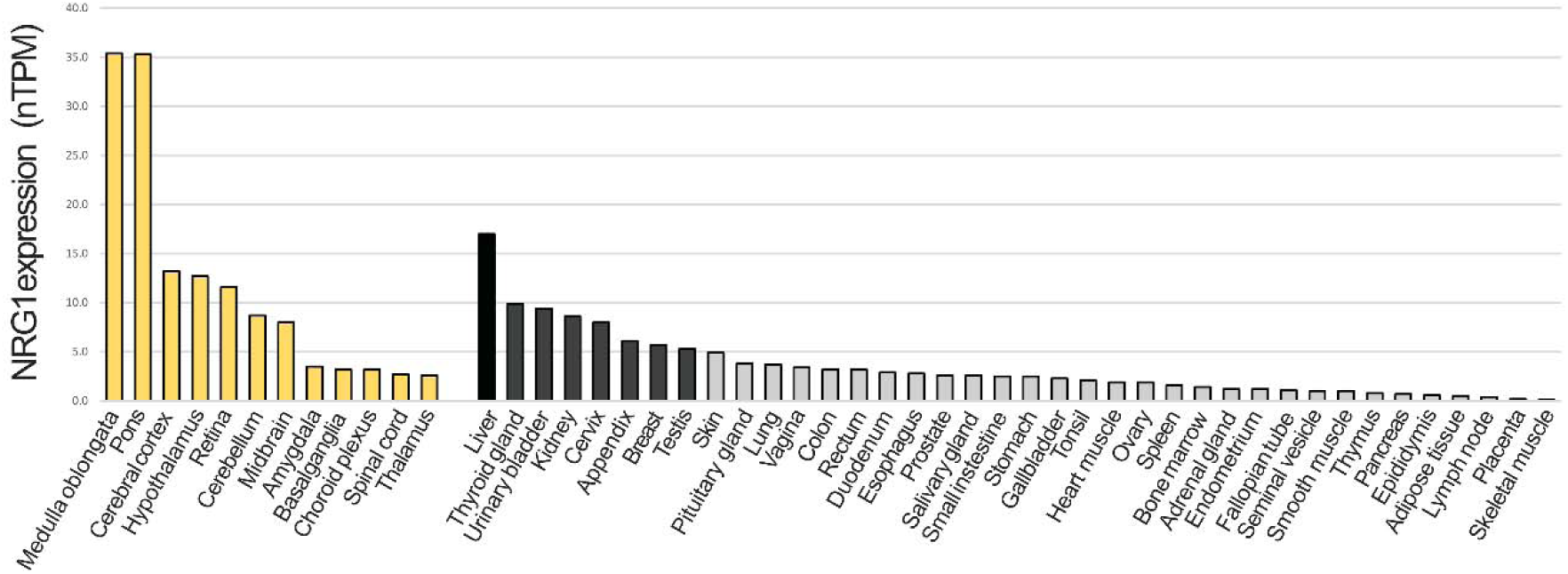
NRG1 expression levels in macrophages presen in a wide variety of human tissues. Quantification of NRG1 mRNA expression in macrophages present different human tissues according to The Human Protein Atlas (Karlsson et al., 2021). The y axis shows the number of transcripts per million. Yellow bars = neural tissue; Grey scale bars = levels of expression in all other tissues.

**Supplementary Figure 5.**
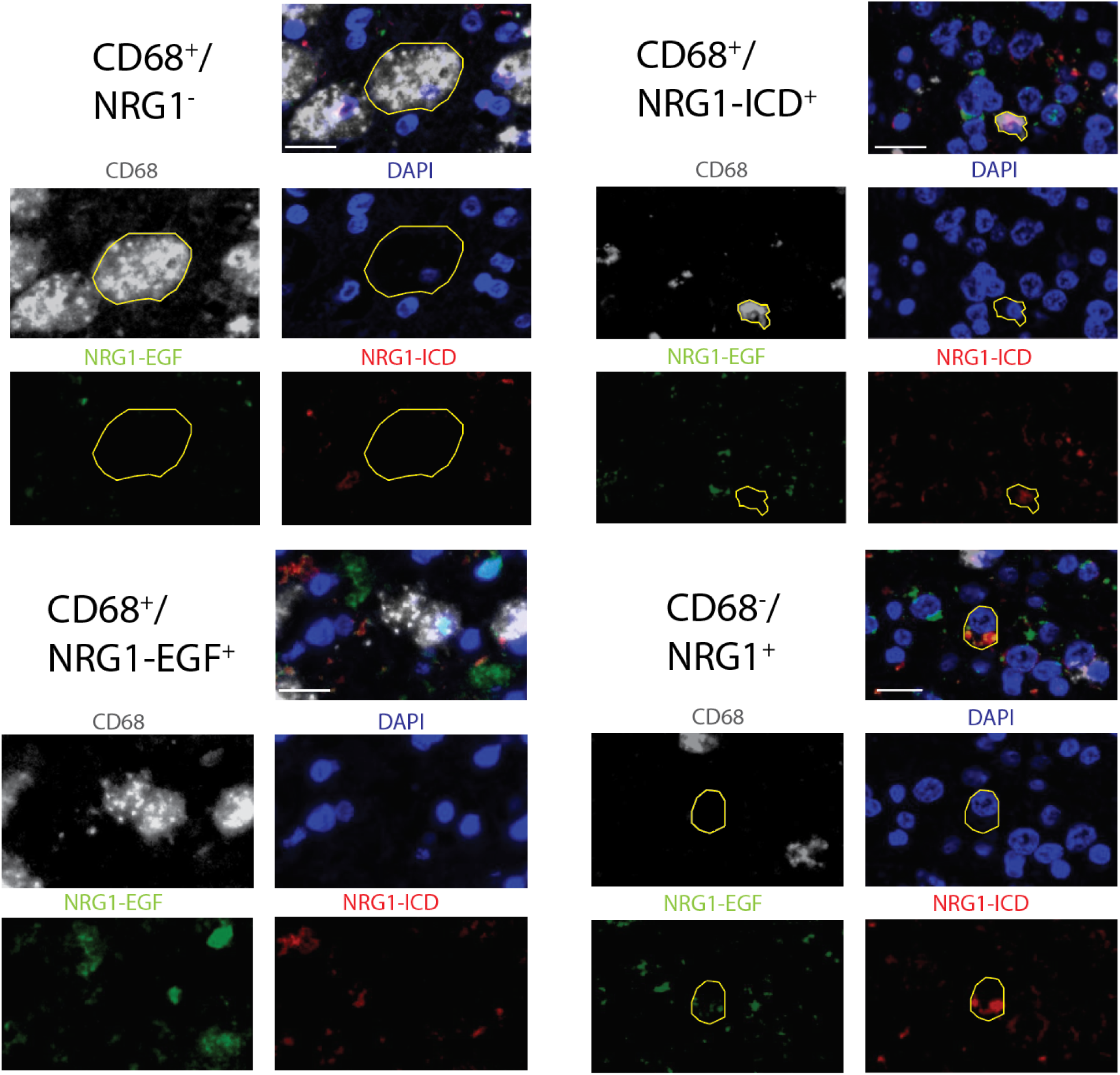
Panel of marker combinations present in GBM samples. CD68^+^ cells are diverse in their NRG1 expression patterns. Panels show CD68^+^ cells that can be NRG1^-^, or positive for either of the NRG1 antibodies independently. CD68^-^ cells were also observed to be positive for presence NRG1. Scale bars = 20 μm.

## Availability of data and materials

FAST5 and BAM files are available from ENA under accession PRJEB62796 (https://www.ebi.ac.uk/ena/browser/home). The different NRG1-VII isoform sequences and expected proteins are available at NCBI under submission numbers OQ272754-OQ272776 (https://www.ncbi.nlm.nih.gov/). The processed data and scripts used in this study are available at https://github.com/wellslab/NRG1-VII_Amplicons. IsoLamp is available on github at https://github.com/ClarkLaboratory/IsoLamp.

## Authorship contributions

Conceptualization MABR, CAW, Data curation MABR, RDP, YDJP, Formal analysis MABR, RDP, YDJP, Funding acquisition CAW, MBC, Investigation MABR, CAW, Methodology MABR, RDP, YDJP, MD, SSW, JG, WD, NR, AL, JH, Project administration CAW, Resources CAW, MBC, TM, Software MABR, RDP, YDJP, JG Supervision CAW, MBC, Validation MABR, RDP, YDJP, MD, SSW, Visualization MABR, MD, SSW, writing original draft MABR, Writing – review and editing MABR, RDP, YDJP, MD, SSW, WD, NR, AL, TM, MBC, CAW.

## Acknowledgements.

The authors thank Professor Matt Watt (Department of Anatomy and Physiology, University of Melbourne), for gifting adipose tissue samples. The authors thanks Dr Le Christy Ying and Prof Paul Hertzog (Hudson Institute, Clayton, Melbourne) for gifting 2D macrophage RNA for this study. The authors thank Mrs Zahra Elahi (Department of Anatomy and Physiology, University of Melbourne) for her logistic and emotional support, crucial to this study. The authors thank Mr Yidi Deng (Department of Maths and Statistics, University of Melbourne) for discussions about statistical approaches.

## Sequence Accessions

All mRNA sequences have been deposited at NCBI GenBank nucleotide database, Accessions assigned 18/1/23: OQ272754, OQ272755, OQ272756, OQ272757, OQ272758, OQ272759, OQ272760, OQ272761, OQ272762, OQ272763, OQ272764, OQ272765, OQ272766, OQ272767, OQ272768, OQ272769, OQ272770, OQ272771, OQ272772, OQ272773, OQ272774, OQ272775, OQ272776.

